# Hoverfly responses to looming stimuli depend on elevation and speed

**DOI:** 10.64898/2026.05.11.724450

**Authors:** Aika H. Young, Jaxon Mitchell, Katja Sporar Klinge, Andrew B. Barron, Yuri Ogawa, Karin Nordström

## Abstract

An object on immediate collision course generates a rapidly expanding visual stimulus on the retina, which will typically trigger a fast reaction, such as an evasive behavior. In hoverflies, for example, visual looming stimuli may be generated if the insect is about to collide with a stationary object in the surround, by an approaching predator, or by conspecifics during territorial interactions. Supporting these behavioral responses is a diverse range of looming sensitive descending neurons that project information from the optic lobes and central brain to the motor control centers in the thoracic ganglia. We here show that the looming sensitive descending neurons are predominantly sensitive to looming stimuli located in the ventral visual field. To investigate if this is matched by behavior, we recorded how tethered hoverflies responded to looming stimuli presented dorsally or ventrally on a visual monitor, at four different speeds (*l/|v|* of 10 - 667 ms), covering a naturalistic range. We found that ventral stimuli, especially at intermediate speeds (*l/|v|* = 50 or 200 ms), triggered much stronger behavioral responses than dorsally displayed stimuli. The behavioral data thus not only match the receptive fields of the neurons likely to support the behavior, but also highlight that behavioral output is not entirely reflexive but is strongly modulated by stimulus speed and elevation.

**Significance Statement:** If someone throws a ball at you, this generates a rapidly expanding object across your visual field, which will make you react before you have even had time to think. You may for example duck, dip or dive to avoid the ball, or bring your hands up to grab it. Similarly, many insects respond to rapidly approaching objects. We here show that hoverfly reactions to such looming stimuli depend on stimulus speed and elevation, with the strongest response to stimuli approaching from below. We further demonstrate that the neurons likely supporting these behaviors show highest sensitivity in the ventral visual field, suggesting a close match between neural tuning and behavioral output.

## Introduction

Effectively responding to a rapidly approaching stimulus is crucial to the survival of many animals, as it could indicate an imminent collision with a stationary obstacle or an approaching predator. For instance, mice may either freeze or hide in response to looming stimuli (Yilmaz and Meister, 2013), as would be ecologically appropriate if approached by a predator. Similarly, zebrafish larvae respond to looming objects with fast escape responses away from the stimulus (Dunn et al., 2016). Furthermore, the approach rate of the looming stimulus affects the behavioral latency (Bhattacharyya et al., 2017), since the zebrafish larvae would have more time to evade if the presumed predator is approaching more slowly. Similarly, invertebrates also need to avoid potential predators. When crabs, for example, are approached by looming stimuli they run to their burrow if the burrow is close, but freeze if it is far away (Oliva et al., 2024), and flying insects attempt to escape approaching threats by changing their flight direction (Muijres et al., 2014).

While looming stimuli are often treated as threatening, there are nuances to this. For example, a looming stimulus triggers an escape response in perching *Drosophila*, but if they are already flying, the same looming stimulus leads to a leg extension, which could be indicative of a landing response (Ache et al., 2019a). Furthermore, in flying *Drosophila*, the response depends on the azimuthal location of the looming stimulus, with lateral stimuli causing evasive saccades, while frontal looming stimulus lead to a leg extension (Tammero and Dickinson, 2002). Like other insects, hoverflies need to effectively evade predators, which are likely to approach their prey rapidly from above (but see e.g. Rossoni et al., 2021). In addition, looming stimuli may be generated by conspecifics during territorial and courtship interactions (Thyselius et al., 2018; Thyselius et al., 2023), or by stationary objects directly in the flight path (e.g. Egelhaaf et al., 2014; Maimon et al., 2008).

Supporting these behavioral reactions are looming sensitive neurons found in the optic lobes (e.g. Béron de Astrada et al., 2013; de Vries and Clandinin, 2012; Oliva and Tomsic, 2014), which project to looming sensitive descending neurons that in turn innervate motor neuropils in the thoracic ganglion (e.g. Ache et al., 2019b; Fotowat et al., 2009; Gabbiani et al., 1999). In the hoverfly, some of these looming sensitive descending neurons are also sensitive to widefield optic flow and to target motion (Nicholas et al., 2020; Nicholas et al., 2023). Such dual sensitivity might be useful during territorial interactions when the conspecific switches from being visualized as a small target when distant, to a looming stimulus, as it gets closer (Thyselius et al., 2018). However, where in the visual field they have highest sensitivity to looming stimuli has not been shown. To investigate this, we here recorded intracellularly from looming sensitive descending neurons and show that they are predominantly sensitive to looming stimuli in the ventral visual field.

To investigate if the ventral neural sensitivity is matched by behavior, we displayed dorsal and ventral looming stimuli at four different speeds (*l/|v|* = 10-667 ms), and measured how this affected head, leg and wing movements of tethered hoverflies. We found that the elevation and speed of the looming stimulus affected behavioral output, with an increased wing beat amplitude to most stimuli, especially at slower speeds, while leg extensions were exclusive to ventral looming stimuli, predominantly at medium speeds (*l/|v|* = 50 and 200 ms). Our data thus show that not only do the receptive fields of looming sensitive descending neurons match behavior, but also that these responses are not purely reflexive. Instead, hoverflies appear to integrate stimulus speed and elevation to generate distinct behavioral outputs.

## Material and Methods

### Animals

*Eristalis tenax* hoverflies were reared and housed as described previously (Nicholas et al., 2018). In short, female *Eristalis tenax* hoverflies were collected under permit from the Wittunga Botanic Garden in Adelaide, South Australia, and their larvae raised at room temperature in a rabbit dung slurry. Adult hoverflies were housed in square bugdorms (24.5 cm side, Australian Entomological Supplies) and kept at a 12 h light:12 h dark cycle. Light was provided by Apollo G2 lights (Pracht Lichttechick GmbH, Germany) with a color temperature of 4000K providing a light intensity of 540 lux. The temperature during the light period was set to 22 ± 1°C and during the dark to 18 ± 1°C. They had unrestricted access to water, sugar, and pollen. The hoverflies used in electrophysiology were 38-128 days old, and in behavior the female hoverflies were 31-78 days old.

### Electrophysiology

Recordings were made from 30 looming sensitive descending neurons in 15 male and 15 female *Eristalis tenax* hoverflies. At experimental time the hoverfly was immobilized ventral side up using a beeswax and resin mixture, and a small incision made at the anterior end of the thorax to expose the cervical connective. The trachea covering the cervical connective were removed using forceps, and the gut partially pulled out if it exhibited excessive movement. If the recording area appeared dry, a small amount of PBS (Sigma-Aldrich) was added to the dissected region. A small wire hook was placed under the cervical connective for mechanical support, and a silver wire used as a reference electrode.

Aluminosilicate electrodes were pulled using a Sutter P-1000 micropipette puller (Sutter Instruments, Novato, California, USA) resulting in resistances of 40 to 70 MΩ. The electrodes were backfilled with 1 M KCl and inserted into the cervical connective using a PM-10 Piezo translator (World Precision Instruments Inc, Sarato, Florida, USA). When a neuron was impaled, a drop in voltage was observed. Only neurons with a stable resting potential and action potential amplitude of at least 10 mV were used. Data were amplified using an Axoclamp-2B amplifier (Axon Instruments, Australia), and 50 Hz noise minimized with a Humbug (Quest Scientific, North Vancouver, BC, Canada). Data acquisition and digitization were performed at 10 kHz using an NI USB-6210 16-bit data acquisition card (National Instruments, Austin, Texas, USA) and the MATLAB Data Acquisition Toolbox.

### Tethered flight

We tethered the anterior end of the hoverfly’s dorsal thorax to a needle (BD Microlance 23G x 1 1/4” - 0.6 x 30 mm Blue hypodermic needles) at a 32° angle (see Nicholas et al., 2026) using a mixture of bee’s wax and resin. The needle was connected to a syringe (BD tuberculin syringe, 1 ml). We provided airflow manually to encourage flight (Ogawa et al., 2025) and once flying consistently, we placed the hoverfly in the center of the tethered flight arena. If the hoverflies stopped flying, we encouraged flight by providing a manual perch followed by quickly removing it to prompt takeoff. In the instances where this did not induce sustained flight, we removed the hoverfly from the arena and provided a sugar-water solution and manual air flow to encourage flight. Hoverflies that still did not attempt to fly at any point during at least one full trial, even when prompted, were removed from the experiment.

We filmed the hoverfly from above at 100 Hz (Kaushik et al., 2020; Ogawa et al., 2025) using a FLIR Blackfly S USB camera (BFS-U3-04S2M-C Mono, Edmund Optics Technician, Barrington, NJ, USA) with a spatial resolution of 720 x 540 pixels. 100 Hz is below the wing beat frequency of *Eristalis* hoverflies (Walker et al., 2010), thus allowing at least one entire wingbeat to be smeared across each frame of the recording. The camera was equipped with an infrared pass filter (R-72, Edmund Optics Technician, Barrington, NJ, USA), the hoverfly illuminated with infrared lights (AL-314IR-940-30B-002, A-Bright Industrial Co Ltd, Taiwan, mounted in JANSJÖ LED USB lamps, IKEA, Sweden), and a musou black surface (Shin Kokushoku Musou black, KOYO Orient Japan, Saitama, Japan) placed below to maximize the contrast (see also Ogawa et al., 2025).

### Visual stimuli

In electrophysiology, the hoverfly was positioned 10 cm away from the center of the linearized monitor (ViewSonic, California, USA), which had a refresh rate of 165 Hz and a spatial resolution of 2560 (width) x 1440 (height) pixels, covering 144° of the fly’s visual field in azimuth and 120° in elevation (Fig. 1A). In behavior, the hoverfly was positioned 11.9 cm away from the center of the linearized monitor (Asus, Beitou District, Taipei, Taiwan), which had a refresh rate of 165 Hz and a spatial resolution of 1440 (width) x 2560 (height) pixels, covering 109° of the fly’s visual field in azimuth and 137° in elevation (Fig. 2A).

**Figure 1.**
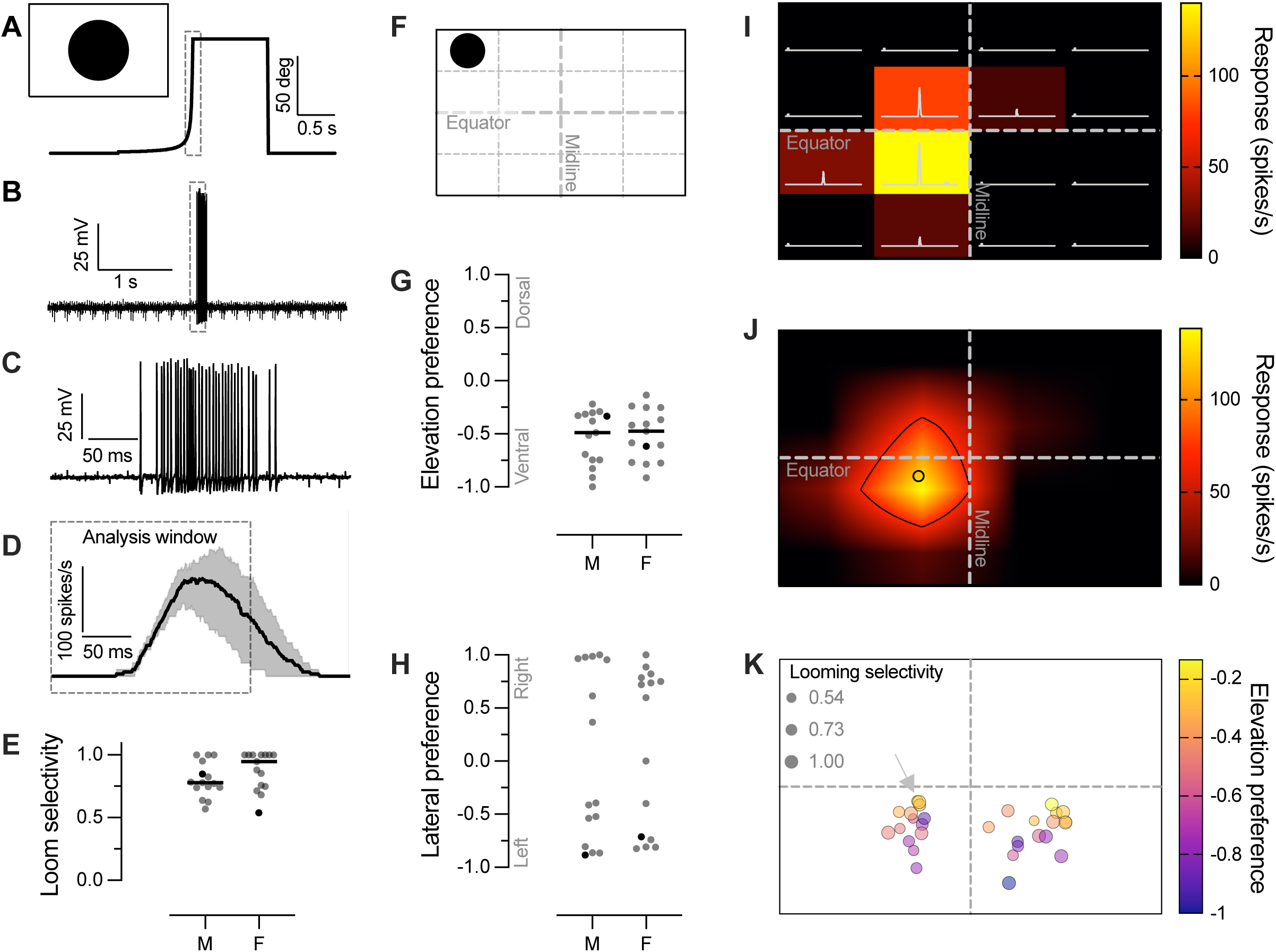
Looming neurons have ventral receptive fields. A) The inset pictogram displays the maximum size (diameter = 96°) and relative location of the looming stimulus used in electrophysiology. The growth rate of the looming stimulus is also shown, where the circular, black stimulus grew for 1 s, then remained stationary at maximum size on the screen for 1 s. Pre- and post-stimulation the screen was white. B) The intracellular response of a female looming sensitive descending neuron, time aligned to the stimulus in panel A. C) Magnification of the response in panel B. D) The mean and range of the response of the same neuron across three repetitions, time-aligned to the data in panel C. In panels A, B, D, the dashed box shows the 200 ms analysis window centered on the time the looming stimulus reached maximum size, used to calculate the average spike rate. E) The looming selectivity across 15 male and 15 female neurons, with the horizontal lines indicating the median. There was not significant difference between males and females (Mann-Whitney test, p = 0.17). F) The location of the smaller looming stimuli used to map receptive fields, shown at maximum size (diameter = 40°). G) The elevation preference across the same 15 male and 15 female neurons, with the horizontal lines indicating the median. There was not significant difference between males and females (Welch’s t-test, p = 0.72). G) The lateral preference across the 15 male and 15 female neurons. I) The response of one example male neuron to the receptive field technique shown in panel F. J) The same data as in panel I, spatially interpolated to higher resolution. The black line shows the 50% response contour, and the black open circle the center of this contour line. K) The receptive field centers across the same 30 neurons as in panels E, G, H, color coded according to elevation preference, with size indicating looming selectivity. The arrow highlights the neuron shown in panels I, J. In panels E, G and H the black male data point highlights the neuron shown in panels I, J, and the black female data point the neuron shown in panels A-D.

**Figure 2.**
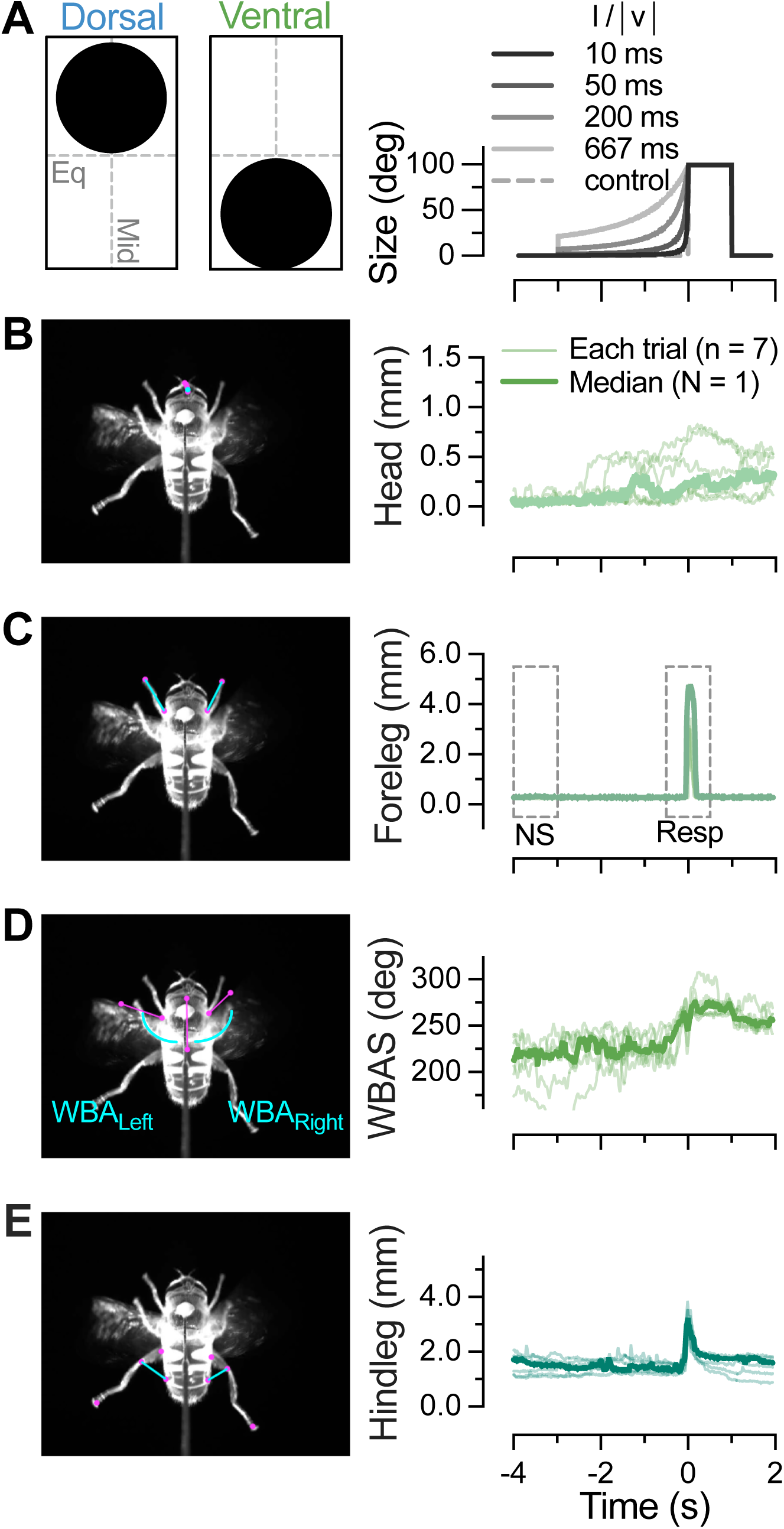
Looming stimulus and body part tracking. A) The pictogram displays the relative location of the dorsal and ventral looming stimuli at maximum size (diameter = 99°) on the left. On the right, the growth rate of the four looming stimuli. In all cases the circular, black stimulus grew for 3 s, then remained stationary at maximum size on the screen for 1 s. As the final size and growth time was the same, each stimulus had a different size at appearance. Pre- and post-stimulation the screen was white. B) The response to the ventral looming stimulus with an *l/|v|* of 200 ms where the magenta dots shows the points on the head that were tracked using DeepLabCut (Kane et al., 2020), and the cyan dot the one used for the data on the right. This data shows the movement of the head relative to its starting point, across 7 repetitions (thin lines), with the thicker line indicating the median across repetitions in this one example hoverfly. C) The magenta dots track the foreleg tarsal claws and the most proximal part of the forelegs seen in the dorsal view, and the cyan line the vector between these. The data on the right shows the average of the left and right foreleg vector from the same example hoverfly. The dashed boxes indicate 1 s pre-stimulus (no stimulus condition, “NS”), and 1 s centered on the time the looming stimulus reached maximum size (response, “Resp”). D) By tracking two points on each wing, and two points along the longitudinal axis we could extract the wing beat amplitude (cyan) of the left (WBA_Left_) and right (WBA_Right_) wing. The data on the right shows the sum of the WBA_Left_ and WBA_Right_. E) The hindleg vector was defined as the distance between middle of the lateral abdomen and the joint between the tibia and femur (cyan line). The data on the right shows the average of the left and the right hindleg vector across the 7 repetitions.

We used custom written scripts (https://github.com/HoverflyLab/FlyFly/releases/tag/v4.2.3) based on the Psychophysics toolbox (Brainard, 1997; Pelli, 1997) in MATLAB (Mathworks) to generate the visual stimuli. We generated looming stimuli using an *l/|v|* of 10 ms for electrophysiology (Fotowat and Gabbiani, 2007; Nicholas et al., 2020). This resulted in a stimulus where a small black circle (1°) appeared in the center of the white monitor and expanded to a final diameter (96°) over 1 s. It remained in full size on the monitor for 1 s. A motion-free control was a black circle with diameter of 96° appearing and remaining on the screen for 2 s. To map the looming receptive field, the visual monitor was divided into a 4×4 grid. Small looming stimuli (*l/|v|* = 10 ms, 1° to 40° expansion over 1 s, remaining on the screen for 1 s, inter stimulus interval of 1 s) were presented at randomized locations within the grid.

For behavior, we generated black looming stimuli on a white background on the dorsal or ventral half of the screen (Fig. 2A). The stimuli were preceded by a minimum 1 s pre-stimulus and 1 s post-stimulus times, during which the screen was white. When looming, the black disc grew for 3 s, using four different *l/|v|* (10 ms - 667 ms). The final size (always the same, 28 cm diameter, 99°) remained on the screen for 1 s before disappearing (Fig. 2A). A motion-free control was a black circle with a diameter of 99° appearing and remaining on the screen for 1 s. In between each presentation, we showed a refresher starfield stimulus (Leibbrandt et al., 2021) sideslipping at 50 cm/s for 10 s to maintain consistent flight. The order of looming stimuli was randomized, but kept constant across flies, and repeated until each hoverfly stopped flying.

### Electrophysiology data analysis

All quantification was programmatically done in MATLAB (R2024b, Mathworks, USA) to minimize data analysis bias. To generate spike histograms, we applied a 50 ms square-wave filter with a resolution of 0.1 ms. The mean response for each neuron was then identified using an analysis window of 100 ms either side of the looming stimulus first reaching maximum size (dashed box, Fig. 1A, B, D). Looming-sensitive neurons were identified based on a stronger response to the looming stimulus compared with the motion-free control (Nicholas et al., 2020) by calculating looming selectivity (Fig. 1E): *(Resp_Loom_ − Resp_Control_) / (Resp_Loom_ + Resp_Control_)*. We kept data from all neurons that had a looming selectivity above 0.25, using a previously published threshold (see Nicholas et al., 2020).

As the hoverfly was placed ventral side up in electrophysiology experiments, we rotated the receptive fields 180° to display them dorsal side up (Fig. 1F, I-K). For each location within the 4x4 grid (Fig. 1F), we extracted the response histogram (Fig. 1I) and calculated the mean spiking frequency within the same 200 ms analysis window as above (dashed box, Fig. 1A, B, D) and indicated the response magnitude with color coding (Fig. 1I). For each neuron, we calculated the elevation preference, by comparing the responses to the eight locations above the equator with the eight locations below the equator: *(Resp_Dorsal_ – Resp_Ventral_) / (Resp_Dorsal_ + Resp_Ventral_)*. We calculated the lateral preference, by comparing the responses to the eight locations to the right of the midline, with the eight locations to the left: *(Resp_Right_ – Resp_Left_) / (Resp_Right_ + Resp_Left_)*.

For each neuron we spatially interpolated the responses to a 200x200 matrix (color coding, Fig. 1J). We quantified the 50% response maximum using MATLAB’s *contour* function (black outline, Fig. 1J). We created a polygon-shape of this 50% contour line using MATLAB’s *polyshape* function and used the *centroid* function to identify the center of the polyshape (open circle, Fig. 1J).

### Behavioral analysis

We used DeepLabCut (DLC) version 2.3.3 (Kane et al., 2020; Nath et al., 2019) to train a model to track 24 points on the hoverfly (magenta dots, Fig. 2B-E, supp movie 1-10). Four DLC models were trained individually, to track the forelegs, head, wings and hindlegs, respectively. For the forelegs, 70 video frames were extracted and manually labelled from videos of two individual animals. To train the head, wing and hindleg models, 45, 46 and 47 videos frames were extracted and labelled respectively, from videos of five individual animals.

To minimize experimenter bias, all quantification was done programmatically in MATLAB (R2024b, Mathworks, USA). For the head, we had tracked 6 points along the midline and measured the deviation of the second most posterior point (cyan, Fig. 2B) from its average position during the 1 s immediately preceding visual stimulation. For the forelegs, we quantified the length of the foreleg vector, defined as the distance between the tarsal claws and the most proximal part of the femur as seen in the dorsal view (cyan lines, Fig. 2C). We used the mean of the left and right foreleg vector (Fig. S1C, D) for further analysis. For the wings, we used two points along the animal’s longitudinal axis, and two each along the anterior edge of each wing stroke. We used this to calculate the wing beat amplitude, defined as the angle between the wing edge and the longitudinal axis (WBA, cyan, Fig. 2D; see e.g. Maimon et al., 2010). We used the sum (WBAS, Fig. 2D) of the left and the right WBA (WBA_Left_+ WBA_Right_; Fig. S1E, F) and the WBA difference (WBAD = |WBA_Left_ - WBA_Right_|) for further analysis. Before calculating the WBAD, WBA_Left_ and WBA_Right_ were normalized to their average position during the 1 s immediately preceding visual stimulation. For each hindleg we labelled the tarsal claw, the joint between the tibia and femur, and the most proximal part of the femur seen in the dorsal view, as well as the lateral sides of the mid-abdomen. We calculated the vector between the abdominal point and the joint between the tibia and femur, and used the mean of this vector from the right and the left hindleg for further analysis (cyan line, Fig. 2E, Fig. S1G, H).

We kept data from all trials where the hoverfly flew continuously from the start of the refresher sideslip stimulus to the end of the subsequent looming stimulus, and for all stimuli that had been repeated at least twice in each animal. For each animal, we quantified the median response across repetitions (thick lines, right, Fig. 2B-E) for further analysis. We used a 1 s analysis window immediately preceding stimulus onset (“NS”, Fig. 2C) and one centered on the time of the looming stimulus first reaching maximum size (“Resp”, Fig. 2C).

Within these we identified the maximum response. To investigate leg extensions further, we calculated an occurrence ratio, defined as the ratio of trials where the foreleg and hindleg vectors were both above 2.5 mm (dashed lines, Fig. 5B). This threshold was based on data from two animals, where we manually identified three frames from nine movies showing leg extension responses to ventral stimuli at the three faster speeds (top, Fig. 5A). We compared this to data from the same two flies, where we manually identified three frames each from nine movies showing no leg extension responses to dorsal or ventral stimuli at any speed (bottom, Fig. 5A).

For leg extensions above 2.5 mm we determined the time of maximum response to ventral stimuli. We used the fact that the time of the response, where time is the time relative to collision (TOC), is directly correlated with the *l/|v|* (Gabbiani et al., 1999), following:

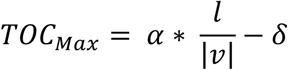

Where α is the slope and δ the intercept. We could then extract the threshold angle (θ_Threshold_) that triggers the behavior using:

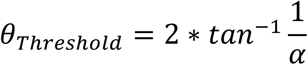

Thus, θ_Threshold_ gives us the angular threshold that triggers a response (here, maximum leg extension) with a delay of δ (Gabbiani et al., 1999).

### Statistical analysis

Statistical analysis was performed in Prism (GraphPad Prism version 11.0.0). The sample size, statistical test, and *p*-values are indicated throughout the text, where *n* refers to the number of repetitions within one animal, and *N* to the number of animals. In all figures, statistical significance is indicated by asterisks, with a single asterisk (*) for p < 0.05, double asterisks (**) for p < 0.01, triple (***) representing p < 0.001, and **** for p < 0.0001.

### Data availability statement

All data and analysis scripts used are openly available in DataDryad: https://doi.org/10.5061/dryad.nk98sf84j. Software used for displaying stimuli and acquiring data can be accessed via GitHub (https://github.com/HoverflyLab/FlyFly/releases/tag/v4.2.3; https://github.com/HoverflyLab/SampSamp; https://github.com/HoverflyLab/FlyFlyDataMerger).

## Results

### Looming-sensitive descending neurons respond more strongly to looming stimuli in the ventral visual field

To characterize looming-sensitive neurons, we performed intracellular recordings in the cervical connective while visually stimulating hoverflies (male and female *Eristalis tenax*). We presented a black looming stimulus in the center of a white screen, with an *l/|v|* of 10 ms and a final size of 96° (Fig. 1A). Looming neurons are defined by their strong responses to looming stimuli (Fig. 1B-D) and small or no response to the appearance control (Nicholas et al., 2020; Nicholas et al., 2023). We calculated the mean across repetitions in each neuron (black, dashed box, Fig. 1D) and compared this to the response to a motion-free control, to calculate the looming selectivity (Fig. 1E). Neurons with a looming selectivity above 0.25 are defined as looming neurons (Nicholas et al., 2020).

We mapped the receptive field using smaller looming stimuli (40° diameter) presented in random order within a 4x4 matrix across the visual monitor (Fig. 1F). The example data from a single male neuron show the response at each location (histograms, Fig. 1I) with the color coding reflecting the mean spiking frequency within the analysis window (as in Fig. 1D). We calculated the elevation preference for each neuron and found that they all had a ventral preference (Fig. 1G), i.e. all neurons responded more strongly to the 40° diameter looming stimuli in the ventral visual field. In contrast, the lateral preference was bimodally distributed (Fig. 1H). There was no significant difference between males and females (Fig. 1E, G, H).

We identified each neuron’s receptive field center (open circle, Fig 1J), and plotted this across neurons, color coded according to elevation preference (Fig. 1K). This analysis highlights that all receptive field centers were located below the visual equator, and that the further away the receptive field center was from the equator, the stronger the elevation preference (color coding, Fig. 1K, R^2^ = 0.60). However, the looming selectivity strength was not correlated with receptive field center location (circle size, Fig. 1K).

### Tethered hoverflies react strongly to looming stimuli

To investigate how tethered hoverflies respond to looming stimuli at different elevations, we used an expanding black disc presented on either the dorsal or ventral part of the monitor (maximum size shown in Fig. 2A). We used four different growth speeds (*l/|v|*, Fig. 2A), ranging from 10 ms (fastest) to 667 ms (slowest). Similar speeds have been used to study neural and behavioral responses to looming stimuli in zebrafish, insects and crabs (see e.g. Ache et al., 2019b; Cámera et al., 2020; Fotowat and Engert, 2023; Fotowat et al., 2009; Fotowat and Gabbiani, 2007; Oliva et al., 2024).

We filmed the tethered hoverflies at 100 Hz from above and used trained DeepLabCut (Kane et al., 2020; Nath et al., 2019) models to track the head, fore- and hindlegs, and wings. The data from an example hoverfly show that when a ventral looming stimulus approaches with an *l/|v|* of 200 ms, the different body parts move substantially, especially close to the time when the looming stimulus reaches its maximum size (time = 0, Fig. 2, supp movie 9). While the head movement is more subtle (Fig. 2B), both forelegs extend rapidly and substantially (Fig. S1A-D, Fig. 2C), the wing beat amplitude increases (Fig. S1E, D; WBAS, Fig. 2D), and the hindlegs extend away from the abdomen (Fig. S1G, H, Fig. 2E). For each animal, we calculated the median across trials (thick lines, right, Fig. 2B-E) and used this to identify the maximum response within a 1 s analysis window (“Resp”, Fig. 2C). As control, we quantified the maximum response with no stimulation (“NS”, Fig. 2C).

### The response depends on stimulus speed and location

To visualize the dependence on stimulus speed and elevation, we plotted the data from each body part as a function of time, color coded according to location (grey = control, blue = dorsal, green = ventral, Fig. 3, N = 9, supp movie 1-10). The data suggest that all body parts move the most to looming stimuli placed in the ventral part of the monitor, especially when these expand at the middle speeds (central green columns, Fig. 3). We quantified the maximum response from each hoverfly within the analysis windows (dashed boxes, Fig. 2C) and confirmed that head movement depends on stimulus elevation (dorsal vs ventral, p = 0.0479, 2-way ANOVA), and stimulus speed (p = 0.0033, Fig. 4A). The head movement was larger than the no-stimulus control to most ventral stimuli (green data, Fig. 4A).

**Figure 3.**
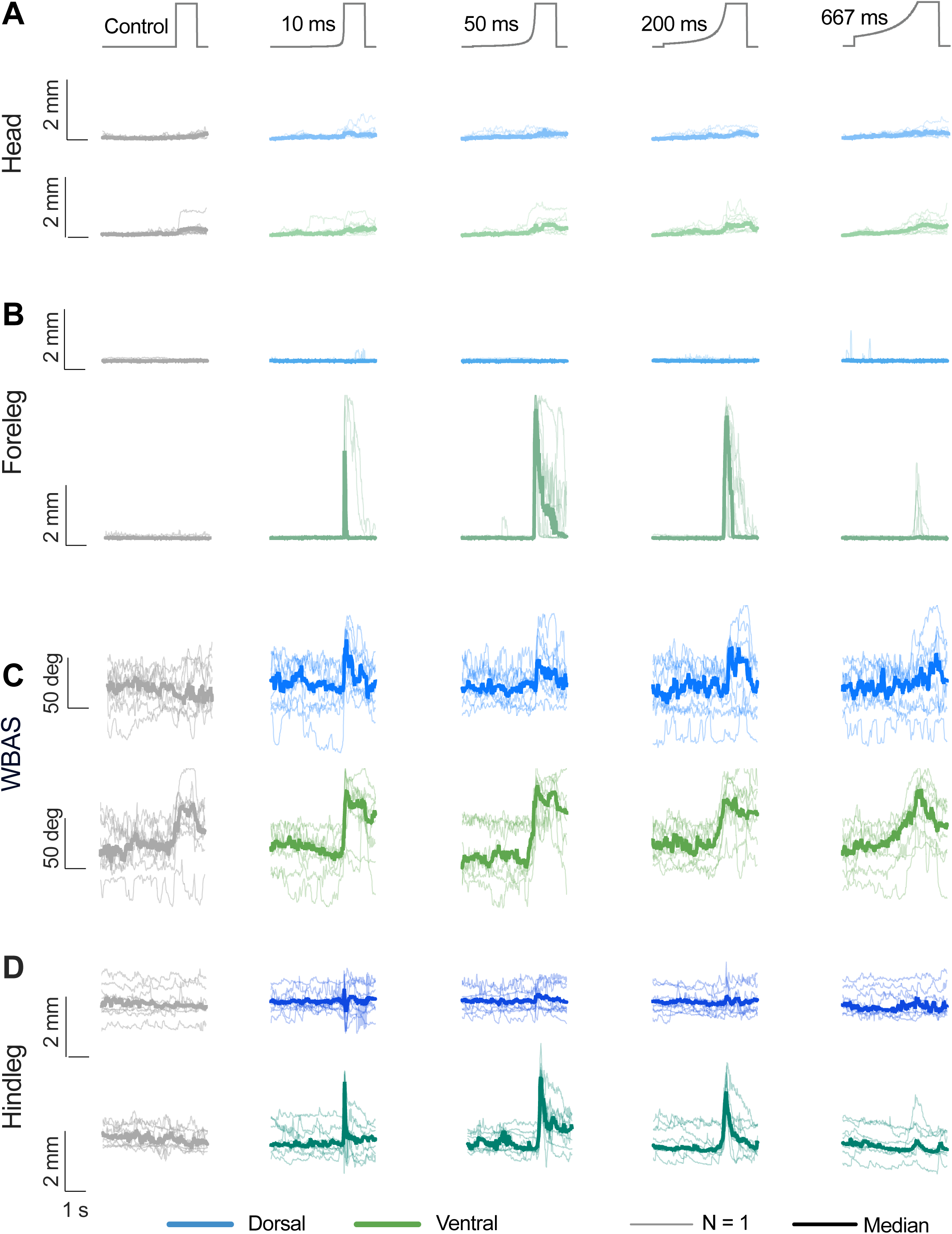
Ventral stimuli at middle speeds generate the largest response. A) *Top.* The stationary control, and looming stimuli at four different speeds. *Middle*. The head movement to dorsal looming stimuli. *Bottom*. The head movement to ventral stimulation. B) The foreleg vector in response to looming stimuli placed in the dorsal (blue) or ventral (green) part of the monitor. C) The WBAS in response to looming stimuli placed in the dorsal (blue) or ventral (green) part of the monitor. D) The hindleg vector in response to dorsal (blue) or ventral (green) looming stimuli. In all panels, the thin lines indicate the median response from each hoverfly, and the thick line the median across flies (N = 9).

**Figure 4.**
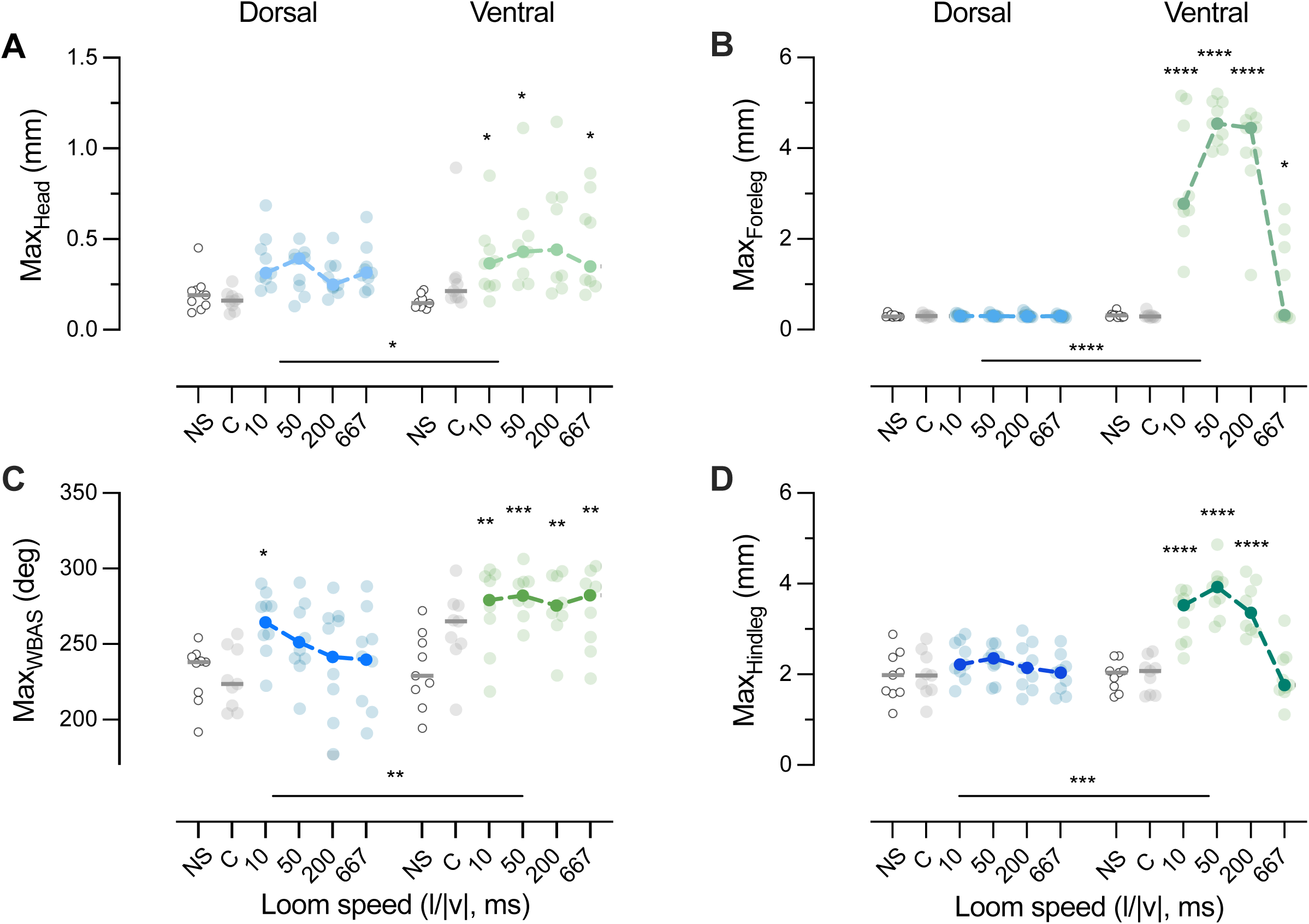
Ventral stimuli generate significantly larger responses. A) Maximum head movement across all flies as a function of the speed of the looming stimulus. The transparent dots show the response from each fly, and the more salient symbols indicate the median across flies. B) Maximum foreleg extension across flies. C) Maximum WBAS across flies. D) Maximum hindleg extension across flies. In all panels, the open symbols show the no-stimulus data, the grey symbols show the responses to the stationary controls, the blue data responses to dorsal stimuli and the green data the responses to ventral stimuli. Statistical significance was investigated using 2-way ANOVA (N = 9) with one star * for p < 0.05, ** for p < 0.01, *** for p < 0.001 and **** for p < 0.0001 compared with the no-stimulus control.

The forelegs did not extend when viewing dorsal looming stimuli (blue data, Fig. 3B, supp movie 2-5), nor the control stimuli (grey data, Fig. 3B, supp movie 1, 6). There was therefore a strong dependence on both stimulus speed and elevation (both p < 0.0001, 2-way ANOVA, Fig. 4B), with a much larger foreleg extension to fast ventral looming stimuli (green data, Fig. 4B, supp movie 7-9) compared with no stimulus (NS, open symbols, Fig. 4B).

In contrast, the wing beat amplitude sum (WBAS) increased to all four looming speeds, at both stimulus elevations, as well as to the ventral appearance control (grey, Fig. 3C), even if the reactions appeared to be stronger to ventral stimuli (green, Fig. 3C). Indeed, the maximum WBAS depended on both stimulus speed (p < 0.0001, 2-way ANOVA) and elevation (p = 0.0032, Fig. 4C), but the variation across animals was large. The dorsal looming stimuli only generated a significant response compared with no stimulus at its fastest growth speed (*l/|v|* = 10 ms, blue data, Fig. 4C).

While the WBAS suggests that the flies could be trying to fly faster, or change altitude (e.g. Zanker, 1990), we next asked if they were trying to fly away from the looming stimulus by quantifying the wing beat amplitude difference (WBAD). This is often used as an indication of yaw turning (e.g. Götz, 1968; Maimon et al., 2010; Tammero et al., 2004). As the looming stimuli were placed along the visual midline, both left and right turns indicate evasive turns. To mitigate this ambiguity, we quantified the absolute WBAD, and found that it increased to all four looming speeds, at both stimulus elevations, as well as to the ventral appearance controls (Fig. S2A-C). Indeed, the increased WBAD compared with no stimulus was significant for several looming speeds, but there was no dependence on elevation (p < 0.0001 for stimulus speed, p = 0.5943 for elevation, 2-way ANOVA, Fig. S2D).

Like the forelegs (Fig. 3A), the hindlegs did not move substantially in response to either the appearance controls (grey data, Fig. 3D), nor to dorsal looming stimuli (blue, Fig. 3D), but the response was strong to ventral stimuli, especially at the three faster speeds (green, Fig. 3D, 4D). The maximum hindleg extension depended strongly on both stimulus speed (p < 0.0001, 2-way ANOVA, Fig. 4D) and elevation (p = 0.003), with significantly stronger responses to ventral stimuli compared with no stimulus conditions at the three faster speeds (green data, Fig. 4D).

### Leg extension responses to ventral stimuli

The data above show similarities between the fore- and hindleg’s dependence on stimulus speed and elevation (Fig. 3B, D, 4B, D). To investigate if these movements are synchronized, we manually identified 27 frames where we could see clear extensions of all six legs (top, Fig. 5A) and compared these with 27 frames without similar leg extensions (bottom, Fig. 5A). By plotting the hindleg extension as a function of the foreleg extension, we find that they are correlated (red data, Fig. 5B) and larger than during other behaviors (grey data, Fig. 5B). We also found that in these frames the midleg extension was correlated with the forelegs (Fig. 5C) and the hindlegs (Fig. 5D).

**Figure 5.**
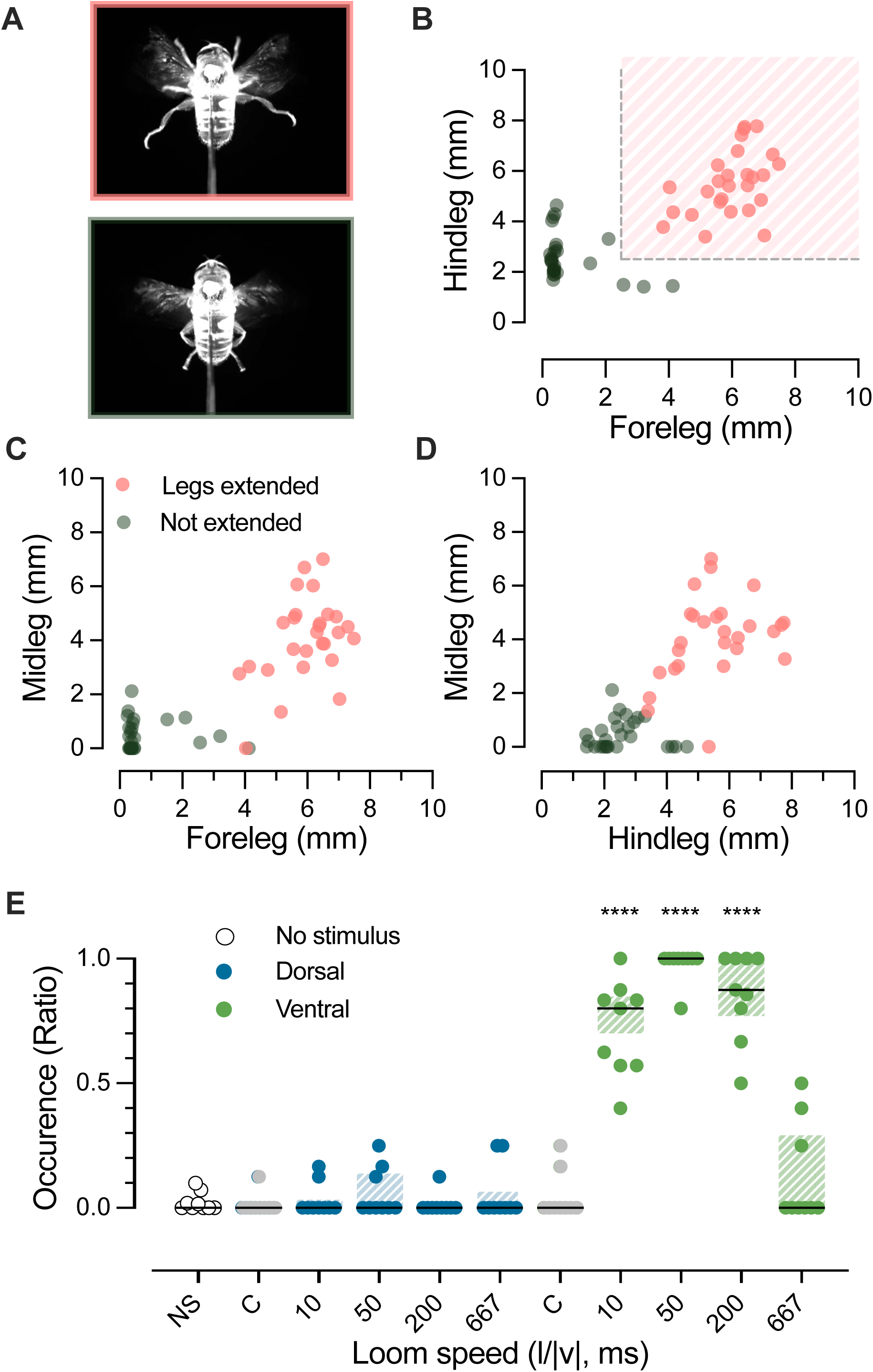
Landing responses depend strongly on stimulus location. A) A typical example of a leg extension (*Top*), and an example frame with no leg extension (*Bottom*). B) Hindleg extension as a function of the foreleg extension in frames that were manually identified to represent typical leg extension (n = 27 frames, from N = 2 animals; red) or not (n = 27 frames, from the same N = 2 animals; grey). The dashed lines indicate the thresholds used to programmatically identify leg extensions. C) Midleg extension as a function of the foreleg extension color coded as in panel B. D) Midleg extension as a function of hindleg extension. E) The occurrence of leg extensions as a function of the speed of the looming stimulus when the stimulus was dorsal (blue) or ventral (green). The open symbols show the no-stimulus data (NS), the grey symbols show the responses to the stationary controls (C). The dots show the occurrence ratio for each fly, the black lines the median across hoverflies, and the shaded areas the interquartile ranges (N = 9). Statistical significance was investigated using 2-way ANOVA (N = 9) with four stars (****) for p < 0.0001 compared with the no-stimulus control.

We used thresholds based on the manually identified leg extensions (dashed lines, red shading, Fig. 5B) and quantified the ratio of leg extensions within a 1 s window surrounding the time of the stimulus reaching maximum size (time = 0, colored shading, Fig. S3) across repetitions in each animal. We compared this to the ratio of leg extensions that occurred before stimulus presentation (grey shading, Fig. S3). This analysis confirms that control and dorsal stimuli did not trigger leg extensions (medians = 0, grey and blue data, Fig. 5E) nor were leg extensions induced without a stimulus (open symbols, NS, Fig. 5E). However, in response to ventral stimuli, leg extensions were induced in ratios of 0.80, 1.0 and 0.88 for the 10 ms, 50 ms and 200 ms stimulus speeds (p < 0.001, 2-way ANOVA, green data, Fig. 5E), but not for the slowest stimulus (*l/|v|* = 667 ms; median = 0, Fig. 5E).

### Leg extension timing

To investigate this leg extension further, we looked closer at the timing of the maximum response to ventral stimuli. The four looming stimuli approach at different speeds, and therefore have different projected time-of-collision with the viewer (Gabbiani et al., 1999) relative to the time it reached maximum size (TOC, Fig. 6A). For example, the fastest stimulus is projected to collide with the viewer 8 ms after reaching maximum size, whereas the slowest stimulus reaches maximum size 567 ms before time-of-collision (dashed lines, Fig. 6A). We can use this information in a threshold model (Gabbiani et al., 1999), which assumes that looming responses are triggered at a fixed delay (δ) after the looming stimulus reaches an angular size threshold (θ_Threshold_) on the retina.

**Figure 6.**
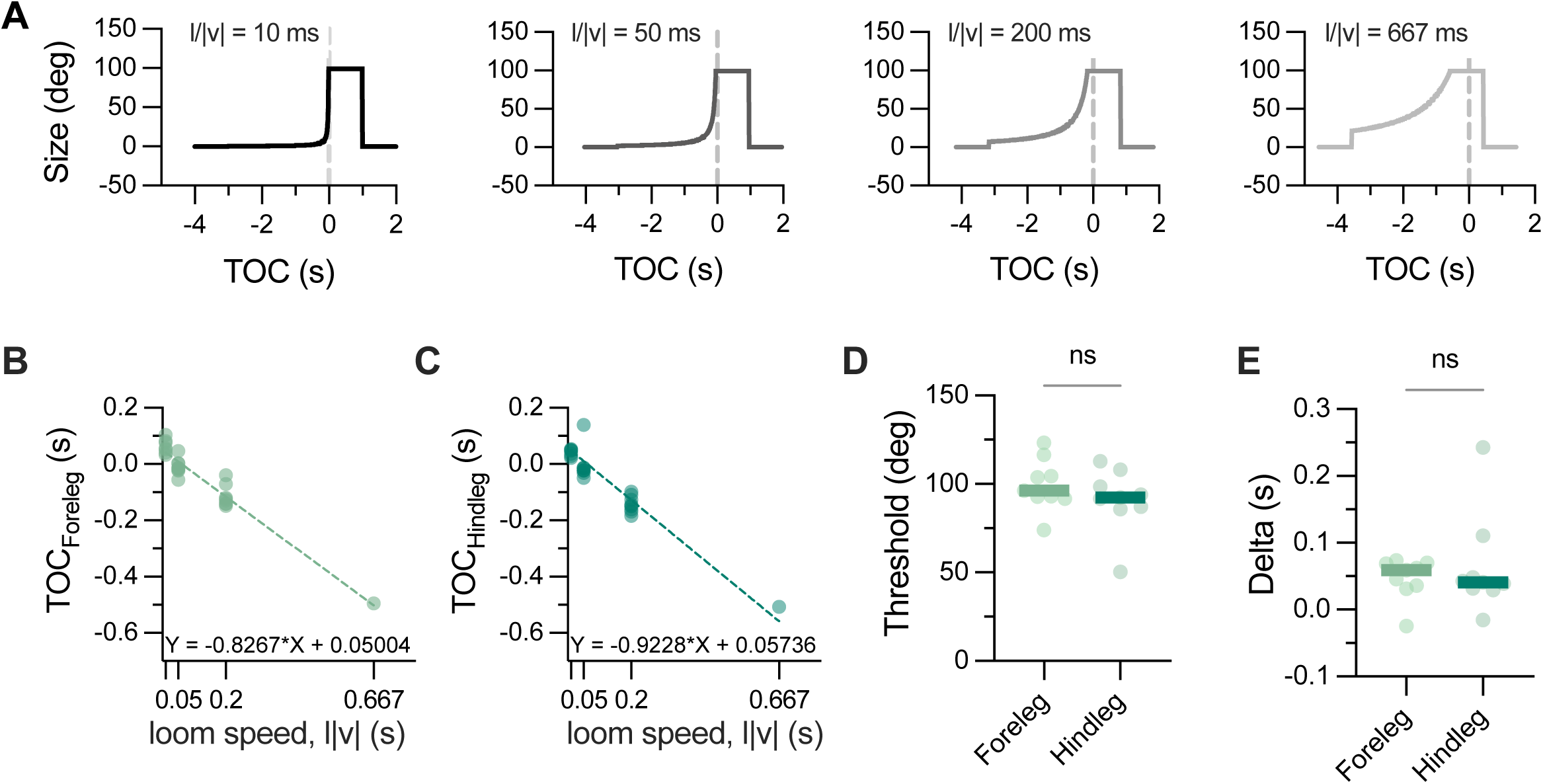
The response time decreases with looming speed. A) The four looming stimuli plotted as a function of projected time-of-collision (TOC), where the dashed lines highlight TOC = 0. B) The time of maximum response of the forelegs as a function of looming speed, for ventral stimuli. The dashed line shows the linear regression to the data across flies that extended their legs > 2.5 mm (N = 7, 9, 8 and 1), with the resulting equation listed at the bottom of the graph. C) The timing relative to TOC for the hindleg extension as a function of looming speed, for ventral stimuli. The dashed line shows the linear regression to the data across flies that extended their legs > 2.5 mm (N = 8, 9, 9 and 1). D) The thresholds extracted from the slope of the linear regression line for each fly (N = 9 and 9). D) The delay extracted from the intercept of the linear regression line for each fly (N = 9 and 9). In panels D and E, statistical significance was investigated using paired t-tests.

This model leads to a linear relationship between the time relative to collision (TOC) of the maximum response and the looming speed (*l/|v|*, Fig. 6B, C), where the slope of the resulting linear regression line is related to the angular size (Gabbiani et al., 1999) and the intercept gives us the delay. Across animals we found that the threshold was 96 ± 49° for the forelegs and 92 ± 62° for the hindlegs (median ± range, Fig. 6D), with no significant difference between the two (p = 0.3196, paired t-test). The delay was 59 ± 9.8 ms for the forelegs and 41 ± 258 ms for the hindlegs (median ± range, Fig. 6E), with no significant difference between the two (p = 0.5517, paired t-test).

## Discussion

We have here shown that looming sensitive descending neurons are predominantly sensitive to looming stimuli in the ventral visual field (Fig. 1). Matching this ventral sensitivity, we found that in tethered hoverflies ventral stimuli generated a strong leg extension (Fig. 2, S1, supp movie 7-10), especially at the middle speeds (green data, Fig. 3, 4). The extensions of the fore- and hindlegs were correlated (Fig. 5, S3) and triggered by similar angular thresholds (Fig. 6). While head movement and WBA increased to both ventral and dorsal stimuli, the response was stronger to ventral stimuli (Fig. 3, 4, S2).

Responding quickly to looming stimuli can be a matter of survival, as such objects may represent predators, or stationary objects in the immediate flight path of the observer. As such, some looming sensitive descending neurons constitute classic examples of command neurons, with wide axons, and stereotyped behavioral output when activated. For example, the fly giant fiber (GF) and locust descending contralateral movement detector (DCMD) trigger fast escape maneuvers as would be ecologically appropriate (e.g. Fotowat et al., 2009; Gabbiani et al., 1999; Gray et al., 2010; Rind and Simmons, 1998). However, while these neurons have been investigated in detail, there are many different invertebrate looming sensitive neurons, that for example vary in their sensitivity and responsiveness to stimulus location and speed (Medan et al., 2015; Namiki et al., 2022; Zacarias et al., 2018). Interestingly, looming sensitivity is also often combined with sensitivity to other visual stimuli, such as widefield or target motion (Olberg, 1986; Rind and Simmons, 1992; Rosner and Homberg, 2013), and some of these neurons are highly multimodal, with e.g. input from mechanoreceptors on the legs and antennae (Ache et al., 2019a).

In hoverflies, there is a diverse group of looming sensitive descending neurons that also respond to widefield and target motion (Nicholas et al., 2023). In these neurons, widefield motion sensitivity is largely located in the ventral visual field, whereas target selectivity is more dorsal (Nicholas et al., 2023). We here showed that looming sensitivity is also predominantly located in the ventral visual field (Fig. 1G, I-K). This separation of sensitivity could for example serve a role during courtship interactions, when the target would initially be small and more dorsal when far away, but be more ventral and looming towards the observer during the later stages of interaction (Thyselius et al., 2018).

Interestingly, looming neurons are presumed to play an important role in predator avoidance, and we would expect at least some predators to approach dorsally. Therefore, the ventral neural sensitivity does not seem to be well matched to behavioral requirements. It is possible that our intracellular electrophysiological technique biased our recordings towards neurons with larger axons, and that there are other looming sensitive neurons with smaller axons and more dorsal receptive fields. Indeed, as the wing beat amplitude increased to the fastest dorsal stimulus (blue data, Fig. 3C, 4C, S2D), this suggests that other descending neurons must have higher dorsal sensitivity, and that these could control evasive maneuvers from fast flying, approaching predators. Indeed, hoverflies are predated upon by avian insectivores, in particular passerines (Hlaváček et al., 2025), and if we assume that a ‘typical’ bird has a wing span of 16 cm, and that it approaches at 10 m/s (Wang et al., 2020), this would have a looming speed, *l/|v|*, of 8 ms (0.08 m / 10 m/s), i.e. close to our fastest speed.

We found that the ventral sensitivity of the looming neurons (Fig. 1) were well matched to stronger behavioral responses to stimuli approaching from below (Fig. 3, 4), especially of the legs. Leg extensions (Fig. 5, 6E) have traditionally been interpreted as a landing response (Ache et al., 2019a), and could indicate that the hoverflies interpreted ventral looming stimuli as a potential flower to land on. Indeed, an ‘average’ flower visited by a hoverfly could have a diameter of 5 cm (see e.g. Dinkel and Lunau, 2001; Gilbert, 1983; Klecka et al., 2018; Lucas et al., 2018; Mishra et al., 2025; Nordström et al., 2017), and if the hoverfly approaches this flower at 0.3 m/s (0.32 m/s: Golding et al., 2001; 0.34 m/s: Thyselius et al., 2018; 0.35 m/s: Thyselius et al., 2023), when the hoverfly approaches the flower would have a looming speed of 83 ms (0.025 m / 0.3 m/s). We found that leg extensions were triggered most by the intermediate speeds (*l/|v|* = 50 and 200 ms, green data, Fig. 3 - 5), thus fitting this ecology well.

However, we found that when the legs extended the wing beat amplitude also increased (green data, Fig. 3, 4). Wing beat amplitude sum has traditionally been used as an indication of flight speed (Nachtigall and Roth, 1983), so this seems counterintuitive as flight speed is reduced as bees approach flowers (Srinivasan et al., 1996). This argues against the hoverflies trying to land on the approaching object, but could instead suggest that they were trying to get away (increased WBAD, Fig. S2, and increased WBAS, Fig. 3C, 4C), but when this did not work due to the open loop nature of our set-up, they were bracing for impact. Indeed, the open loop nature meant that the looming stimuli kept approaching the hoverflies, irrespective of their attempted evasive maneuvers. In contrast, in natural and closed loop systems, behavior shapes what is seen (Ogawa et al., 2025).

Nevertheless, we previously showed that freely flying hoverflies often approached larger beads (4 cm diameter) slowly, followed by landing (Thyselius et al., 2023). When approaching these larger beads, the hoverflies did not change their flight speed as they got closer, instead approaching them at a constant speed (average 0.25 m/s). This argues for the leg extensions being landing responses, and shows that hoverflies do not necessarily reduce their flight speed as they approach an object for landing. Another important consideration when interpreting WBA is that flight speed is affected by many other factors, including the wing frequency, the stroke plane of the wings, wing angles relative to the body, as well as body posture relative to the ground (e.g. Ellington, 1999; Walker et al., 2010). Considering that hoverflies can hover, fly upwards, sideways and even backwards (Geurten et al., 2010), it becomes clear that WBAS alone is a crude metric of flight speed.

We additionally asked whether the hoverflies were trying to turn away from the looming stimulus, by calculating the WBAD (Fig. S2). As the stimuli were placed along the midline, either left or right turns could be evasive, and as such we quantified the absolute WBAD and found that this increased to most stimuli. In the future, it would be interesting to quantify the raw WBAD when using more lateral stimuli to determine whether the flies were trying to evade. Nevertheless, we found that the presumed evasive response did not increase linearly with loom speed (Fig. S2D) as in other species, where e.g. evasive turns in zebrafish are much faster and stronger to faster looming stimuli (Bhattacharyya et al., 2017). Furthermore, in *Drosophila*, there is an inverse relationship between WBAD and landing responses (Tammero and Dickinson, 2002). While we saw no such inverse correlation between WBAD and leg extensions in our data (compare e.g. Fig. 3 and Fig. S2), the WBA increased to dorsal stimuli (blue data, Fig. 3C, 4C, S2), while the legs did not move at all (blue data, Fig. 3B, D, 4B, D). Taken together, this suggests that the pathways underlying leg extensions and collision-avoidance, while distinct, are not as clearly separated in hoverflies as they are in *Drosophila* (Tammero and Dickinson, 2002).

There are, however, some important differences to note. In the *Drosophila* study (Tammero and Dickinson, 2002) the looming stimuli were placed along the visual equator, and the head was fixed relative to the thorax (Lehmann and Dickinson, 1997). In our set-up, the head was free to move (Fig. 2B, 3A, 4A, supp movie 1-10), which could potentially have affected behavioral responses. Both optomotor responses and bar fixation behaviors are strongly affected by the head’s ability to move during tethered flight (see e.g. Cellini and Mongeau, 2020; Fox and Frye, 2014). This suggests that the ability of the head and wings to operate in open vs closed loop could significantly affect behavioral output.

As in previous work (Gabbiani et al., 1999), we used the linear relationship between the response and the estimated time-of-collision, to calculate the delay and the angular threshold that triggers leg extension (Fig. 6). In these equations, the response time is in some papers defined as its onset, and in others as its when it reaches maximum (de Vries and Clandinin, 2012; Fotowat and Gabbiani, 2007; von Reyn et al., 2014), like we did here. We found that the leg extensions used a threshold of about 90° and a delay of about 50 ms (Fig. 6B-E). This matches published accounts, where the locust DCMD has a threshold of 16-34° (Gabbiani et al., 1999) or 40-60° (Stott et al., 2018) and a delay of 15-32 ms (Gabbiani et al., 1999). The looming sensitive Foma neurons in the *Drosophila* brain have a threshold of 67° and a delay of about 40 ms (de Vries and Clandinin, 2012). At the behavioral level, locust escape jumps from a perch have a threshold of 55° and a delay of 57 ms (Fotowat and Gabbiani, 2007), whereas giant fiber induced jumps in *Drosophila* can have a short delay (25-30 ms), or a longer delay when the giant fiber is not used (45 ms, von Reyn et al., 2014). However, in flying *Drosophila*, the landing response triggered by looming stimuli is much slower, with a delay around 150 ms (Ache et al., 2019a; Tammero and Dickinson, 2002). The much faster 50 ms delay to leg extensions that we recorded in flying hoverflies (Fig. 6E) could thus be seen as support of this being an attempt to brace for impact, rather than a landing response.

In the future it would be interesting to film hoverflies in high speed while they are performing natural landing behaviors on flowers, vs moving objects, to determine if the kinematics that we have determined here match free flight behavior and ecological contexts. In addition, it would be interesting to study head free vs head fixed animals to determine whether the head movements (supp video 1-10) help the hoverflies evaluate the ecological relevance of approaching objects.

## CRediT author statement

**Aika Young:** conceptualization, methodology, validation, formal analysis, investigation, writing – review & editing, visualization; **Jaxon Mitchell:** methodology, software, formal analysis, data curation, writing – review & editing, visualization; **Katja Sporar Klinge:** methodology, validation, formal analysis, investigation, writing – review & editing; **Andrew Barron:** methodology, validation, writing – review & editing; **Yuri Ogawa:** conceptualization, methodology, validation, writing – review & editing, supervision; **Karin Nordström:** conceptualization, methodology, software, formal analysis, resources, data curation, writing – original draft, visualization, supervision, project administration, funding acquisition.

## Acknowledgements

We thank Biomedical Engineering at SAHLN and the Botanic Gardens of Adelaide for their ongoing support. This research was funded by the US Air Force Office of Scientific Research (AFOSR, FA9550-23-1-0473) and the Australian Research Council (ARC, DP230100006, DP250100698).

## References

Ache, J. M., Namiki, S., Lee, A., Branson, K. and Card, G. M. (2019a). State-dependent decoupling of sensory and motor circuits underlies behavioral flexibility in *Drosophila*. Nat Neurosci 22, 1132–1139.

Ache, J. M., Polsky, J., Alghailani, S., Parekh, R., Breads, P., Peek, M. Y., Bock, D. D., von Reyn, C. R. and Card, G. M. (2019b). Neural basis for looming size and velocity encoding in the *Drosophila* Giant Fiber escape pathway. Curr Biol 29, 1073–1081.e4.

Béron de Astrada, M., Bengochea, M., Sztarker, J., Delorenzi, A. and Tomsic, D. (2013). Behaviourally related neural plasticity in the arthropod optic lobes. Curr Biol 23, 1389–98.

Bhattacharyya, K., McLean, D. L. and MacIver, M. A. (2017). Visual threat assessment and reticulospinal encoding of calibrated responses in larval zebrafish. Curr Biol 27, 2751–2762.e6.

Brainard, D. H. (1997). The Psychophysics toolbox. Spatial Vision 10, 433–436.

Cámera, A., Belluscio, M. A. and Tomsic, D. (2020). Multielectrode recordings from identified neurons involved in visually elicited escape behavior. Front Behav Neurosci 14, 592309.

Cellini, B. and Mongeau, J. M. (2020). Active vision shapes and coordinates flight motor responses in flies. Proc Natl Acad Sci U S A 117, 23085–23095.

de Vries, S. E. and Clandinin, T. R. (2012). Loom-sensitive neurons link computation to action in the *Drosophila* visual system. Curr Biol 22, 353–62.

Dinkel, T. and Lunau, K. (2001). How drone flies (*Eristalis tenax* L., Syrphidae, Diptera) use floral guides to locate food sources. J Insect Physiol 47, 1111–1118.

Dunn, Timothy W., Gebhardt, C., Naumann, Eva A., Riegler, C., Ahrens, Misha B., Engert, F. and Del Bene, F. (2016). Neural circuits underlying visually evoked escapes in larval zebrafish. Neuron 89, 613–628.

Egelhaaf, M., Kern, R. and Lindemann, J. P. (2014). Motion as a source of environmental information: a fresh view on biological motion computation by insect brains. Front Neural Circuits 8.

Ellington, C. P. (1999). The novel aerodynamics of insect flight: applications to micro-air vehicles. Journal of Experimental Biology 202, 3439–3448.

Fotowat, H. and Engert, F. (2023). Neural circuits underlying habituation of visually evoked escape behaviors in larval zebrafish. Elife 12.

Fotowat, H., Fayyazuddin, A., Bellen, H. J. and Gabbiani, F. (2009). A novel neuronal pathway for visually guided escape in *Drosophila melanogaster*. J Neurophysiol 102, 875–85.

Fotowat, H. and Gabbiani, F. (2007). Relationship between the phases of sensory and motor activity during a looming-evoked multistage escape behavior. J Neurosci 27, 10047–59.

Fox, J. L. and Frye, M. A. (2014). Figure-ground discrimination behavior in *Drosophila*. II. Visual influences on head movement behavior. J Exp Biol 217, 570–9.

Gabbiani, F., Krapp, H. G. and Laurent, G. (1999). Computation of object approach by a wide-field, motion-sensitive neuron. J Neurosci 19, 1122–1141.

Geurten, B. R., Kern, R., Braun, E. and Egelhaaf, M. (2010). A syntax of hoverfly flight prototypes. J Exp Biol 213, 2461–75.

Gilbert, F. (1983). The foraging ecology of hoverflies (Diptera, Syphidae): circular movemens on composite flowers. Behav Ecol Sociobiol 13, 253–257.

Golding, Y. C., Ennos, A. R. and Edmunds, M. (2001). Similarity in flight behaviour between the honeybee *Apis mellifera* (Hymenoptera: apidae) and its presumed mimic, the dronefly *Eristalis tenax* (Diptera: syrphidae). J Exp Biol 204, 139–45.

Götz, K. G. (1968). Flight control in *Drosophila* by visual perception of motion. Kybernetik 4, 199–208.

Gray, J. R., Blincow, E. and Robertson, R. M. (2010). A pair of motion-sensitive neurons in the locust encode approaches of a looming object. J Comp Physiol A Neuroethol Sens Neural Behav Physiol 196, 927–38.

Hlaváček, A., Mikula, P., Hadrava, J. and Lučan, R. K. (2025). Migrating hoverflies as potential food source for co-migrating insectivorous birds. R Soc Open Sci 12, 241743.

Kane, G. A., Lopes, G., Saunders, J. L., Mathis, A. and Mathis, M. W. (2020). Real-time, low-latency closed-loop feedback using markerless posture tracking. Elife 9, e61909.

Kaushik, P. K., Renz, M. and Olsson, S. B. (2020). Characterising long-range search behavior in Diptera using complex 3D virtual environments. Proc Natl Acad Sci 117, 12201–12207.

Klecka, J., Hadrava, J., Biella, P. and Akter, A. (2018). Flower visitation by hoverflies (Diptera: Syrphidae) in a temperate plant-pollinator network. PeerJ 6, e6025.

Lehmann, F.-O. and Dickinson, M. H. (1997). The changes in power requirements and muscle efficiency during elevated force production in the fruit fly *Drosophila melanogaster*. J Exp Biol 200, 1133–1143.

Leibbrandt, R., Nicholas, S. and Nordström, K. (2021). The impulse response of optic flow-sensitive descending neurons to roll m-sequences. J Exp Biol 224.

Lucas, A., Bodger, O., Brosi, B. J., Ford, C. R., Forman, D. W., Greig, C., Hegarty, M., Neyland, P. J. and de Vere, N. (2018). Generalisation and specialisation in hoverfly (Syrphidae) grassland pollen transport networks revealed by DNA metabarcoding. J Anim Ecol 87, 1008–1021.

Maimon, G., Straw, A. D. and Dickinson, M. H. (2008). A simple vision-based algorithm for decision making in flying *Drosophila*. Curr Biol 18, 464–70.

Maimon, G., Straw, A. D. and Dickinson, M. H. (2010). Active flight increases the gain of visual motion processing in *Drosophila*. Nat Neurosci 13, 393–9.

Medan, V., Beron De Astrada, M., Scarano, F. and Tomsic, D. (2015). A network of visual motion-sensitive neurons for computing object position in an arthropod. J Neurosci 35, 6654–66.

Mishra, A., Jain, A., Iyer, P. S., Suryanarayanan, A., Nordström, K. and Olsson, S. B. (2025). Innate floral object identification in a solitary pollinator employs a combination of both visual and olfactory cues. Naturwissenschaften 112, 18.

Muijres, F. T., Elzinga, M. J., Melis, J. M. and Dickinson, M. H. (2014). Flies evade looming targets by executing rapid visually directed banked turns. Science 344, 172–177.

Nachtigall, W. and Roth, W. (1983). Correlations between stationary measurable parameters of wing movement and aerodynamic force production in the blowfly (*Calliphora vicina* R.-D.). Journal of comparative physiology 150, 251–260.

Namiki, S., Ros, I. G., Morrow, C., Rowell, W. J., Card, G. M., Korff, W. and Dickinson, M. H. (2022). A population of descending neurons that regulates the flight motor of *Drosophila*. Curr Biol 32, 1189–1196.e6.

Nath, T., Mathis, A., Chen, A. C., Patel, A., Bethge, M. and Mathis, M. W. (2019). Using DeepLabCut for 3D markerless pose estimation across species and behaviors. Nature Protocols 14, 2152–2176.

Nicholas, S., Leibbrandt, R. and Nordström, K. (2020). Visual motion sensitivity in descending neurons in the hoverfly. J Comp Physiol A 206, 149–163.

Nicholas, S., Ogawa, Y. and Nordström, K. (2023). Dual receptive fields underlying target and wide-field motion sensitivity in looming-sensitive descending neurons. eNeuro 10, ENEURO.0188-23.2023.

Nicholas, S., Sporar Klinge, K., Turnbull, L., Moran, A., Young, A., Ogawa, Y. and Nordström, K. (2026). Sexual dimorphism in sensorimotor transformation of optic flow. Elife 15, RP109795.

Nicholas, S., Thyselius, M., Holden, M. and Nordström, K. (2018). Rearing and long-term maintenance of *Eristalis tenax* hoverflies for research studies. JoVE, e57711.

Nordström, K., Dahlbom, J., Pragadheesh, V. S., Ghosh, S., Olsson, A., Dyakova, O., Suresh, S. K. and Olsson, S. B. (2017). In situ modeling of multimodal floral cues attracting wild pollinators across environments. Proc Natl Acad Sci U S A 114, 13218–13223.

Ogawa, Y., Aoukar, R., Leibbrandt, R., Manger, J. S., Bagheri, Z. M., Turnbull, L., Johnston, C., Kaushik, P. K., Mitchell, J., Hemmi, J. M. et al. (2025). Combining Unity with machine vision to create low latency, flexible and simple virtual realities. Methods in Ecology and Evolution 16, 126–144.

Olberg, R. M. (1986). Identified target-selective visual interneurons descending from the dragonfly brain. J Comp Physiol A 159, 827–840.

Oliva, D., Gültig, M., Cámera, A. and Tomsic, D. (2024). Freezing of movements and its correspondence with MLG1 neuron response to looming stimuli in the crab Neohelice. J Exp Biol 227.

Oliva, D. and Tomsic, D. (2014). Computation of object approach by a system of visual motion-sensitive neurons in the crab *Neohelice*. Journal of Neurophysiology 112, 1477–1490.

Pelli, D. G. (1997). The VideoToolbox software for visual psychophysics: Transforming numbers into movies. Spatial Vision 10, 437–442.

Rind, F. C. and Simmons, P. J. (1992). Orthopteran DCMD neuron: A re-evaluation of responses to moving objects. I. Selective responses to approaching objects. J Neurophysiol 68, 1654–1666.

Rind, F. C. and Simmons, P. J. (1998). Local circuit for the computation of object approach by an identified visual neuron in the locust. J Comp Neurol 395, 405–15.

Rosner, R. and Homberg, U. (2013). Widespread sensitivity to looming stimuli and small moving objects in the central complex of an insect brain. J Neurosci 33, 8122–33.

Rossoni, S., Fabian, S. T., Sutton, G. P. and Gonzalez-Bellido, P. T. (2021). Gravity and active acceleration limit the ability of killer flies (*Coenosia attenuata*) to steer towards prey when attacking from above. J R Soc Interface 18, 20210058.

Srinivasan, M. V., Zhang, S. W., Lehrer, M. and Collett, T. S. (1996). Honeybee navigation en route to the goal: Visual flight control and odometry. J Exp Biol 199, 237–244.

Tammero, L. F. and Dickinson, M. H. (2002). Collision-avoidance and landing responses are mediated by separate pathways in the fruit fly, *Drosophila melanogaster*. J Exp Biol 205, 2785–98.

Tammero, L. F., Frye, M. A. and Dickinson, M. H. (2004). Spatial organization of visuomotor reflexes in Drosophila. J Exp Biol 207, 113–22.

Thyselius, M., Gonzalez-Bellido, P. T., Wardill, T. J. and Nordström, K. (2018). Visual approach computation in feeding hoverflies. J Exp Biol 221, jeb177162.

Thyselius, M., Ogawa, Y., Leibbrandt, R., Wardill, T. J., Gonzalez-Bellido, P. T. and Nordström, K. (2023). Hoverfly (*Eristalis tenax*) pursuit of artificial targets. J Exp Biol 226, jeb244895.

von Reyn, C. R., Breads, P., Peek, M. Y., Zheng, G. Z., Williamson, W. R., Yee, A. L., Leonardo, A. and Card, G. M. (2014). A spike-timing mechanism for action selection. Nat Neurosci 17, 962–70.

Walker, S. M., Thomas, A. L. R. and Taylor, G. K. (2010). Deformable wing kinematics in free-flying hoverflies. J Roy Soc Interface 7, 131–142.

Wang, Y., Yin, Y., Ren, Z., Jiang, C., Sun, Y., Li, J., Nabi, G., Wu, Y. and Li, D. (2020). A comparison of flight energetics and kinematics of migratory Brambling and residential Eurasian Tree Sparrow. Avian Research 11, 25.

Yilmaz, M. and Meister, M. (2013). Rapid innate defensive responses of mice to looming visual stimuli. Curr Biol 23, 2011–2015.

Zacarias, R., Namiki, S., Card, G. M., Vasconcelos, M. L. and Moita, M. A. (2018). Speed dependent descending control of freezing behavior in *Drosophila melanogaster*. Nat Commun 9, 3697.

Zanker, J. M. (1990). The Wing Beat of Drosophila-Melanogaster .3. Control. Philosophical Transactions of the Royal Society B-Biological Sciences 327, 45–64.

