## Supplementary material for "Hoverfly responses to looming stimuli depend on elevation and speed": Supp Text

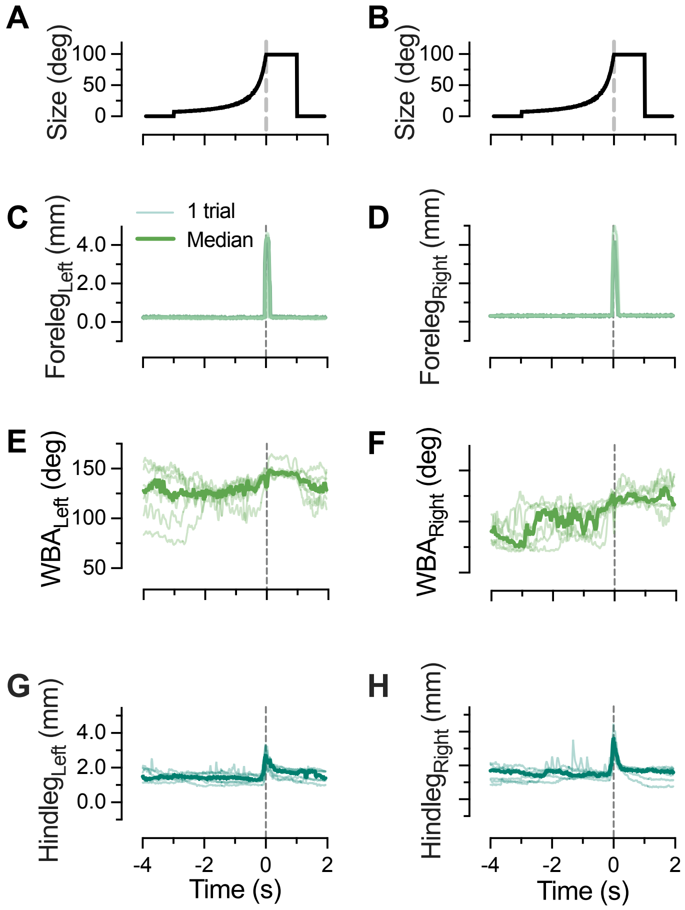


**Supplementary Figure 1. The tracking of body parts.** Associated with Figure 2. A) The looming stimulus with an *l*/|*v*| of 200 ms. B) Same as panel A. C) The left foreleg vector from the same example hoverfly as in Figure 2. D) The right foreleg vector from the same hoverfly. E) The left wing beat amplitude. F) The right wing beat amplitude. G) The left hindleg vector. H) The right hindleg vector. In all panels, the thin lines show the data from individual repetitions (n = 7), and the thick line shows the median from the fly (N = 1).

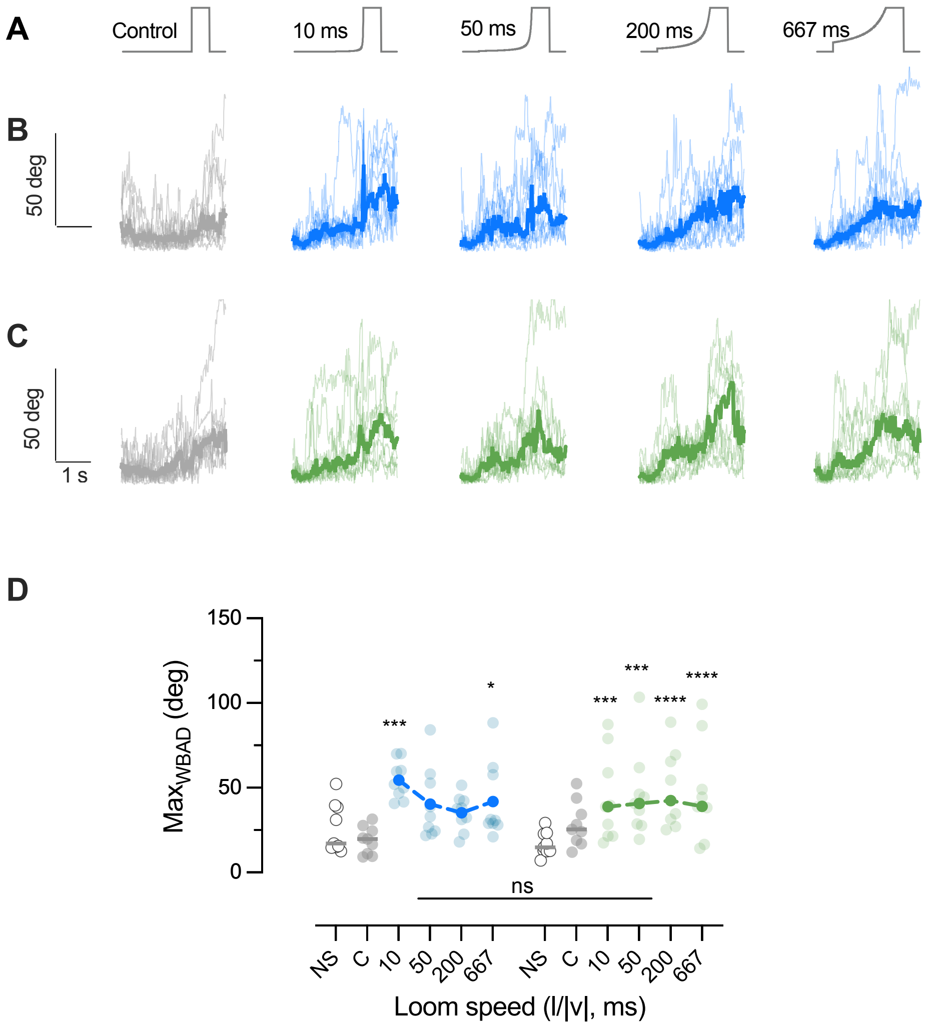


**Supplementary Figure 2. The wing beat amplitude increases to most stimuli.** Associated with Figure 3, 4.

A) The stationary control, and looming stimuli at four different speeds. B) The absolute wing beat amplitude difference (WBAD) in response to dorsal control (grey) and looming stimuli (blue). C) The WBAD in response to ventral control (grey) and looming stimuli (green). In panels B and C, the thin lines indicate the median response from each hoverfly, and the thick line the median across flies (N = 9). D) Maximum WBAD as a function of the speed of the looming stimulus. The transparent dots show the maximum response from each hoverfly, and the more salient symbols indicate the median across flies. The stars indicate significant difference compared with the stationary control (2-way ANOVA, N = 9), with one star * for $p < 0.05$, *** for $p < 0.001$ and **** for $p < 0.0001$.

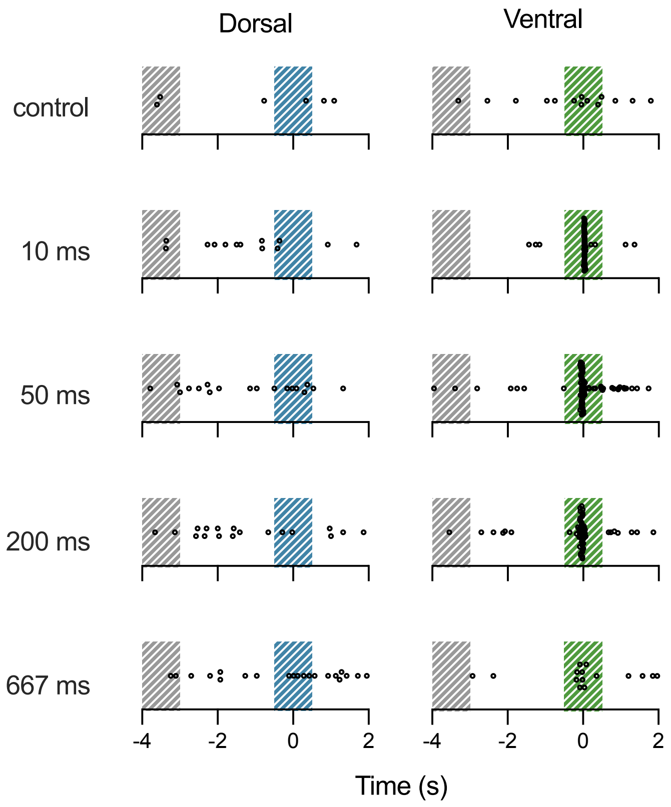


**Supplementary Figure 3. Onset of the leg extension behavior across stimulus conditions and behavioral state.** Associated with Figure 5.
Onset timing for all leg extensions found across all repetitions (n = 2 - 7) in all flies (N = 9). The shaded areas indicate 1 s pre-stimulus (grey) and 1 s surrounding the time of reaching maximum looming size (blue for dorsal, green for ventral stimuli).

### Supplementary Movie legends

**Supplementary Movie 1. Response to dorsal control stimuli.**

The movie shows the response of a tethered hoverfly when responding to a dorsal appearance control. The stimulus on the right is time aligned with the video of the hoverfly. The magenta dots show the automatic tracking done by DeepLabCut (Kane et al., 2020), and the cyan lines the data used for quantification.

**Supplementary Movie 2. Response to dorsal fast stimuli.**

The movie shows the response of a tethered hoverfly when responding to a dorsal looming stimulus with an $l/|v|$ of 10 ms.

**Supplementary Movie 3. Response to dorsal semi fast stimuli.**

The movie shows the response of a tethered hoverfly when responding to a dorsal looming stimulus with an $l/|v|$ of 50 ms.

**Supplementary Movie 4. Response to dorsal semi slow stimuli.**

The movie shows the response of a tethered hoverfly when responding to a dorsal looming stimulus with an $l/|v|$ of 200 ms.

**Supplementary Movie 5. Response to dorsal slow stimuli.**

The movie shows the response of a tethered hoverfly when responding to a dorsal looming stimulus with an $l/|v|$ of 667 ms.

**Supplementary Movie 6. Response to ventral control stimuli.**

The movie shows the response of a tethered hoverfly when responding to a ventral control stimulus.

**Supplementary Movie 7. Response to ventral fast stimuli.**

The movie shows the response of a tethered hoverfly when responding to a ventral looming stimulus with an $l/|v|$ of 10 ms.

**Supplementary Movie 8. Response to ventral semi fast stimuli.**

The movie shows the response of a tethered hoverfly when responding to a ventral looming stimulus with an $l/|v|$ of 50 ms.

**Supplementary Movie 9. Response to ventral semi slow stimuli.**

The movie shows the response of a tethered hoverfly when responding to a ventral looming stimulus with an $l/|v|$ of 200 ms.

**Supplementary Movie 10. Response to ventral slow stimuli.**

The movie shows the response of a tethered hoverfly when responding to a ventral looming stimulus with an $l/|v|$ of 667 ms.
